# Probing Phosphorylation-Induced Vibrational Couplings in CFTR by 2D IR Spectra Simulations

**DOI:** 10.1101/2024.02.27.582059

**Authors:** Jing Zhu, Tunan Chen, Yu Zhao, Guangfu Ji

## Abstract

Phosphorylation of the regulator (R) domain underlies the basis for gating in the human cystic fibrosis transmembrane conductance regulator (CFTR), malfunction or down regulation of CFTR leads to defective apical chloride transport. The biophysical mechanism that underlies the regulatory effect of R domain is still unclear. Here, we utilize a combination of molecular dynamics simulations and theoretically calculated two-dimensional infrared (2D IR) spectra to probe both the structure and spectral signature of phosphorylated and unphosphorylated CFTR. We uncover an ATP-independent asymptotic movement of nucleotide binding domains (NBDs) driven by phosphorylated R domain. Utilizing non-rephasing cross ground-state bleach infrared (GB IR) spectra simulation, we overcome the interpretation hurdle caused by overlaps of multiple vibrational modes, and find distinct vibrational couplings induced by phosphorylation. By calculating exciton eigenfrequencies, we pinpoint specific vibrational couplings to individual amide I modes (carbonyl stretches), unveiling a critical role of serine residues in modulating the coupling state of neighboring amino acids. Our findings offer a bond-specific perspective on how intramolecular interactions within the R domain translate into its broader regulatory function.

## Introduction

CFTR is a chloride channel residing in epithelial tissues, orchestrates the vital secretory functions of various organs and tissues. It governs the flow of ions and water onto the apical surface of epithelia lining our airways, sweat glands, and exocrine organ lumens (1, 2). Malfunctioning or down regulation of CFTR channels trap chloride ions within cells, preventing the crucial hydration of cellular surfaces. This dehydration triggers the formation of thick and viscous mucus that can obstruct airways and cause various symptoms associated with cystic fibrosis (CF) (3, 4), such as chronic sinusitis, nasal polyps, bronchiectasis, primary sclerosing cholangitis, recurrent pancreatitis, and cervical mucus abnormality (5). CFTR belongs to ATP Binding Cassette (ABC) protein family, harboring two transmembrane domains (TMDs) that form the channel pore, each followed by a cytosolic NBD. These two ABC-canonical TMD-NBD halves are linked by the unique cytosolic R domain, an intrinsically disordered peptide segment rich in charged residues and multiple phosphorylation sites (1, 3). The phosphorylated R domain and ATP bound dimerized NBDs of CFTR is coupled to pore opening (3, 6–8), upon ATP hydrolysis, the NBD dimer dissociates and the pore closes (9).

The regulation of CFTR activity can be largely described to an inhibitory influence of the unphosphorylated R domain on channel gating. The direct evidence comes from the electron cryomicroscopy (cryo-EM) unphosphorylated CFTR structure (9, 10). Although the R domain is not well resolved in these structures, its clearly visible density is wedged in between the two NBDs and among the cytosolic extensions of TMDs. It remains unclear how phosphorylation relieves inhibition. If the R domain is completely embedded between the two NBDs, how does cAMP-dependent protein kinase (PKA) reach its target serines? The model suggesting occasional spontaneous release of the unphosphorylated R domain from its occluded position (10) may render it accessible to the kinase, but it is difficult to explain how the R domain overcomes the high energy barriers imposed by the steric hindances from NBDs. Furthermore, the lack of significant chemical shift changes in R domain between nonphosphorylated and phosphorylated CFTR by chemical shift perturbation (CSP) (11) suggests no dramatic local environment transitions. HNCO NMR experiments and R region interaction studies offer alternative insights, that phosphorylation alters the R domain’s binding landscape, reduces its affinity for NBDs (11) but enhances interaction with the N-terminal (aa 46-60) and C-terminal (aa 1438-1480) peptides of CFTR (11–13). Meanwhile, introducing negative charges by substituting phosphorylated Ser sites with aspartate (Asp) (14) elicits a small but substantial activity before exposure to exogenenous PKA, suggesting the electrostatic repulsion may also play a role in the biophysical mechanism that underlines the R domain’s regulatory effects.

The biophysical mechanisms of the regulatory effect of R domain have remained elusive, largely due to the lack of complete structure information of R domain, especially in the field of theoretical research. The complete structures predicted by AlphaFold (15, 16) provide possibilities for research in this filed. In addition, due to the inherent disorder of the R domain in physiological states, Molecular Dynamics (MD) method based on force field derivation of atomic coordinates can fully demonstrate this disorder process on the time scale, which is a naturalistic method for studying disorder structures. Besides, this dynamic process only takes the structure predicted by AlphaFold as the initial value, and over time, it will form more structures that are close to the real state, which to some extent weakens the impact of prediction errors.

2D IR (17–19) provides bond-specific structural resolution and can be applied to resolve reaction pathway with time-scales span from femtoseconds to hours. It has the fast time-resolution to tracking ultrafast processes like electron transfer and solvent dynamics. Cross-peaks (20, 21) are the hallmark of 2D IR spectroscopy, they are a measure of couplings between molecular vibrations depended on their relative distance and orientation. Couplings will cause the vibrational modes to delocalize over large spatial regions, so as to form the characteristic peaks corresponding to the structures, such as the amide I mode (carbonyl stretch: C=O) of random coils (1645 cm^-1^), long α-helices (1650 cm^-1^) and β-sheets (1620 cm-1 and 1680 cm^-1^) (22). Moreover, 2D IR spectra can be quantitatively computed from MD simulations, which solve Newton’s equation of motions for a system of N interacting atoms. Therefore, the calculated 2D IR spectrum is actually a set of vibrations of N interacting oscillators, which provides a theoretical basis for mapping characteristic spectra to individual oscillators.

In this paper, we utilized a combined MD simulation and theoretically calculated 2D IR spectra to investigate the biophysical mechanism that underlies the R domain’s regulatory effect. The MD method captures the dynamic pictures of conformational landscape of the R domain before and after phosphorylation. The non-rephasing cross GB IR spectrum, overcoming the interpretation hurdle caused by overlaps of multiple vibrational modes in the 2D IR spectrum, highlights the differences in the vibrational couplings induced by phosphorylation. By calculating the eigenfrequency of individual amide I modes in the R domain, we assign these couplings to specific oscillator pairs and obtain phosphorylation sites that significantly alter the coupling states. This presents a significant leap forward in our understanding of the R domain’s regulatory mechanism, achieved through bond-specific resolution, for further investigation into the biophysical basis of CFTR related pathologies.

## Results

### 1. MD simulations of the predicted CFTR

Firstly, we compared the sequence differences between the AlphaFold and cryo-EM structures (PDB ID: 5UAK) (10) in Fig. 1A, both of them are in the dephosphorylated and ATP-free form. The cryo-EM structure has 1158 residues with amino acids ranging from 5-402, 439-645, 844-883, 909-1172, 1207-1436, and an unregistered sequence with 19 amino acids. Specifically, in the cryo-EM structure, aa 646-843, which belongs to the R domain, and aa 403-438, which belongs to the regulatory insertion (RI) segment, are almost missing, and all known phosphorylation sites are located in these missing sequences. Exception for those missing sequences, the amino acid sequence of the predicted structure is completely consistent with the cryo-EM structure, with complete aa 1-1480 residues.

**Fig. 1.**
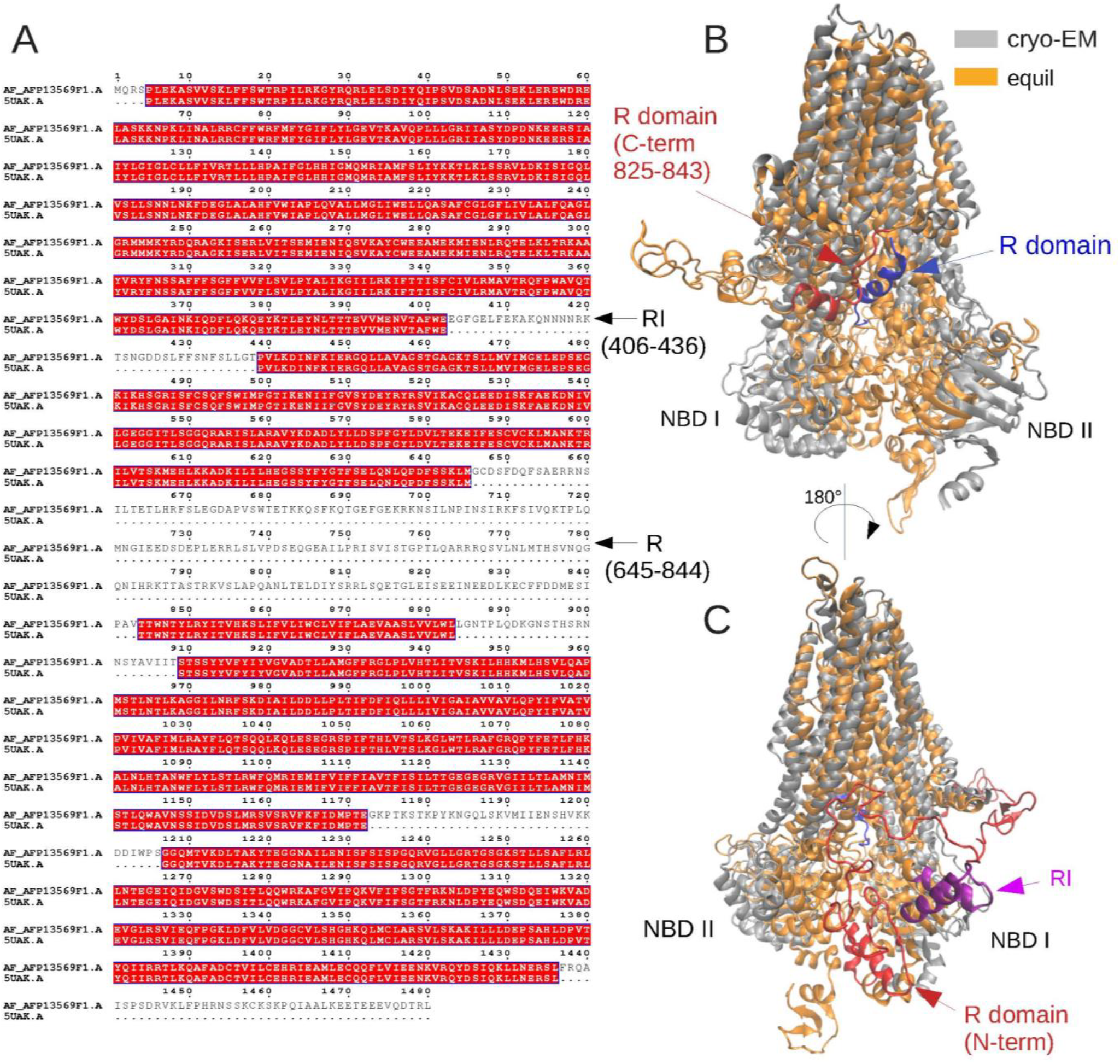
(A) The aligned sequences between AlphaFold and cryo-EM structure. The red highlighted sites represent consistent sequences, the unmarked sites represent sequences deletions in the cryo-EM structure. (B) The 3D structure of MD equilibrium (orange) and (C) cryo-EM (gray). The blue marked structure represents the R domain in cryo-EM structure, red represents that in MD equilibrium structure. The purple marked structure represents the RI domain in MD equilibrium structure.

In Fig. S1, the predicted R domain of CFTR exhibits a dispersed random coil structure, without considering the effects of the lipid film and solvent environment. In order to make this structure closer to the actual situation, we conducted MD simulation in the system (Fig. S1), where the predicted CFTR was embedded in the DOPC membrane and solvated in chlorine ion water. The RMSD results of whole protein and R domain backbones showed that the structure tends to be stable after 50 ns MD simulations (Fig. S2).

The comparison between the 3D structure of the MD equilibrium and the cryo-EM structure is shown in Fig. 1B and C. It indicates that the TMD-NBD halves in the equilibrium structure are closer than those in the cryo-EM structure. Specifically, an α-helical structure (Fig. 1B, colored in blue) that has been resolved by cryo-EM is believed to correspond to aa 825-843 at the C-terminal of R domain (10). These corresponding sequences also forms an α-helix in the MD equilibrium structure (Fig. 1B, colored in red), with a C-terminal orientation similar to that in the cryo-EM structure, both extending towards the interior of the two TMD-NBD halves. The difference is that in the MD equilibrium structure, the N-terminal of the helix is connected to the R domain, forming a chain located on the entire molecule surface. In addition, the N-terminal (aa 648-670) of MD equilibrium structure in the R domain also forms α-helices, they are connected to the NBD I and wedges into the TMD-NBD halves. These helices are relatively flexible, the overall motion of R domain may loose these helices during MD simulation, it seems that these helices store the topological energy for the motion of R domain. β-sheets may also be found in the R domain, but it is a more flexible structure with an unfixed position.

### 2. MD simulation of the phos- and unphos-CFTR

We conducted two sets of experiments according to whether the R domain is phosphorylated or not based on the MD equilibrium structure. The first group simulates the MD of unphosphorylated (unphos-) CFTR without any modification to the equilibrium structure. The second group simulates the MD of phosphorylated (phos-) CFTR, with phosphorylation sites of aa 670, 700, 712, 737, 753, 768, 795 and 813. The equilibrium structures after MD simulation are shown in Fig. 2A and B. Compared to unphos-CFTR, the TMD-NBD halves of the phos-CFTR is closer on the cytoplasmic side. However, there is no significant difference in the secondary structure of their R domains, with the main difference being in their randomly coiled conformation.

**Fig. 2.**
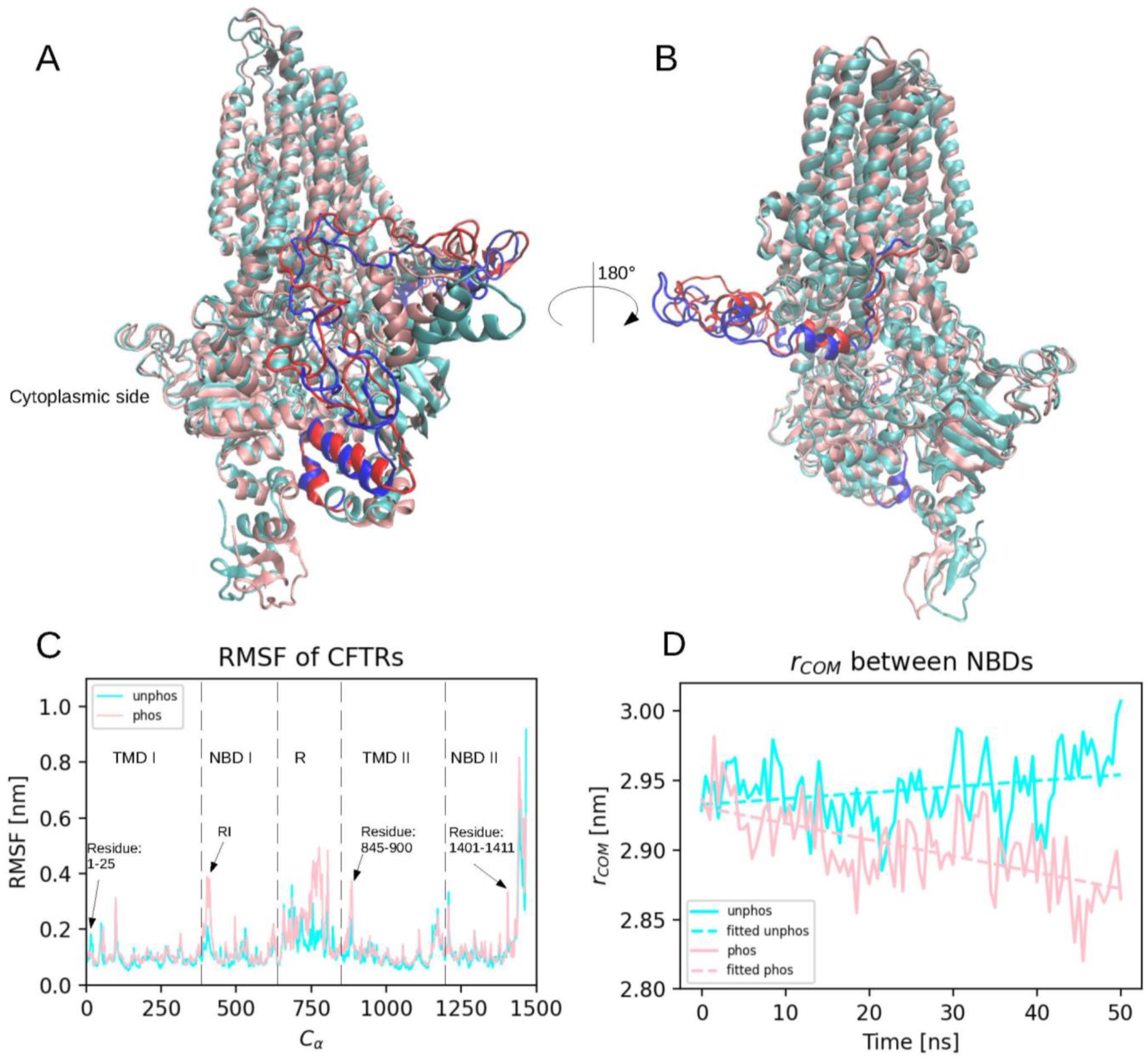
(A and B) The 3D structure of MD equilibrium unphos-CFTR (cyan) and phos-CFTR (pink). The structure marked in blue represents the R domain of unphos-CFTR, red represents the domain of phos-CFTR. (C) The RMSF of Cα position in phos- and unphos-CFTR in the 50 ns trajectory fitted to the initial frame. (D) The distance between two weighted COM of NBDs within 50 ns MD simulation. The dashed lines represent the linear fitting results.

To quantify the fluctuations in the skeleton of R domain, we calculated the root mean square fluctuation (RMSF) of C_α_ positions in the 50 ns trajectory fitted to the initial frame. The result is shown in Fig. 2C, it indicates that the overall fluctuation of C_α_ become more intense after phosphorylation, with particularly significant fluctuations observed in the R domain, the RI segment of NBD1, aa 845-900 of TMD2, and aa 1401-1411 of NBD2.

The fluctuation of skeleton will inevitably cause structural changes, and different fluctuation will shape different structures. To quantify these differences, we calculated the distance over time between the two weighted center of mass (COM) (23) of NBDs. Fig. 2D shows that during 50 ns MD simulation, the NBDs in phos-CFTR gradually approach to each other at a rate ∼ 0.012Å/ns. In unphos-CFTR, the distance of NBDs fluctuates around 2.95 nm, and even slightly increases in the last 10 ns.

### 3. 2D IR spectra of R domain in the unphos- and phos-CFTR

To observe the vibrational differences of amide I modes in the R domain caused by phosphorylation, we conducted another 20 ps MD simulations on the phos- and nuphos-CFTR systems, and recorded the kinetic parameters every 2 fs to calculate the 2D IR spectral of amide I in R domain within 200 vibrators. The spectra are shown in Fig. 3. The FTIR spectrum of unphos-CFTR (Fig. 3A) shows that the absorption peak of amide I modes is at 1680 cm^-1^, with a full width at half-maximum (FWHM) of ∼50 cm^-1^ and a broad shoulder peak at ∼1690 cm^-1^. For phos-CFTR (Fig. 3C), the absorption peak is at 1684 cm^-1^ with a FWHM of ∼50 cm^-1^, and there are three sharp shoulder peaks at 1670 cm^-1^, 1693 cm^-1^, and 1713 cm^-1^, respectively. The MD equilibrium R domain is mainly composed of α-helices, random coils and sometimes β-sheets. Therefore, the expected absorption peak should be around 1650 cm^-1^, which is the characteristic peaks of α-helices and random coils (22). The overall blue shift from 1650 cm^-1^ to 1680 cm^-1^ in our study may be caused by the underestimate of the red-shift by the site-energy parameterization from electrostatic model (Jansen) that occurs upon aqueous solvation of amide I mode from gas phase (24). If we redshift the total spectrum by 30 cm^-1^ to approach the reference value, the two additional shoulder peaks (1640 and 1683 cm^-1^) in phos-CFTR may correspond to the characteristic of β-sheets. In fact, all three secondary structures have characteristic absorption peaks at 1640-1650 cm^-1^, which undoubtedly interferes the ability to distinguish them through FTIR spectroscopy.

**Fig. 3.**
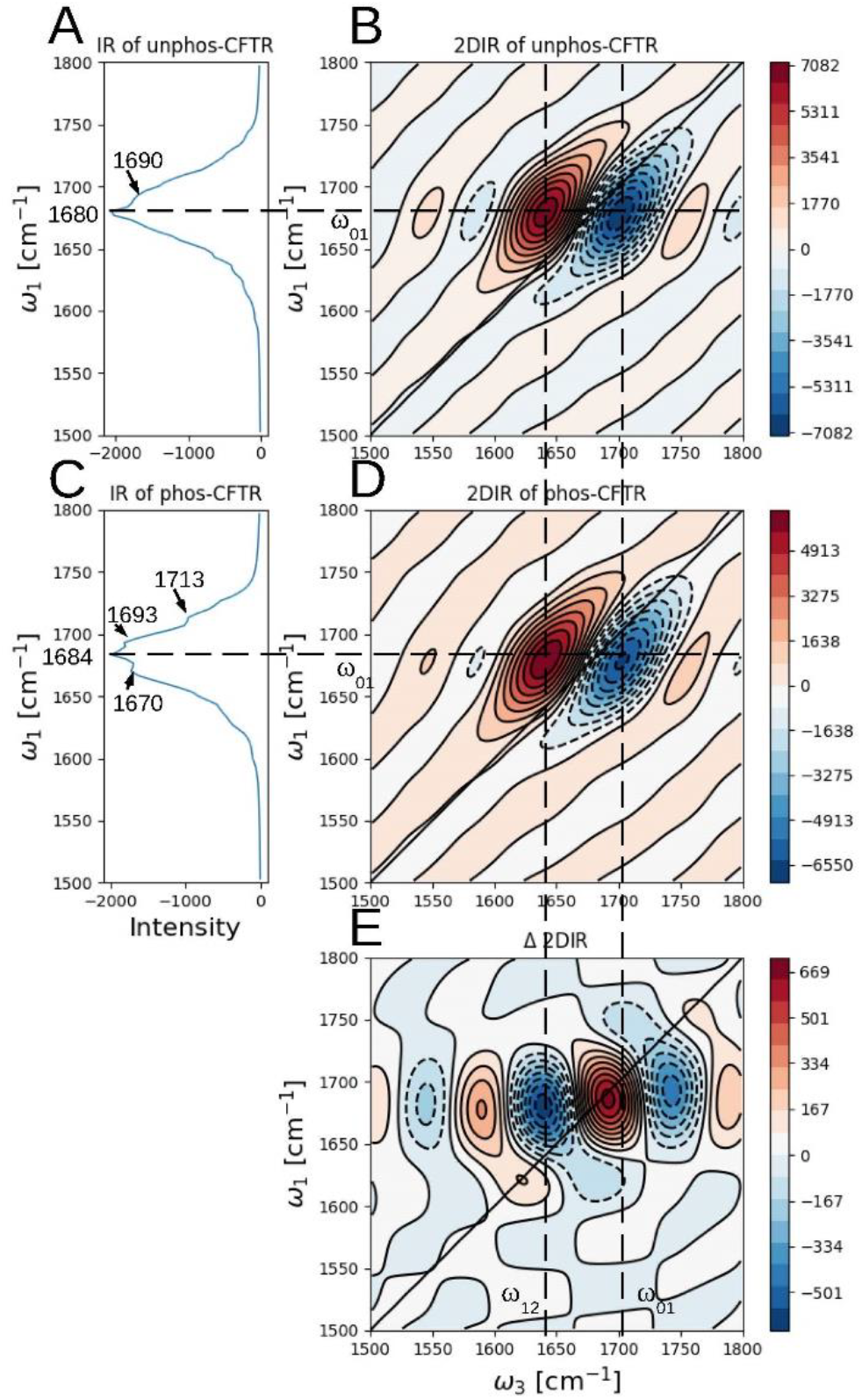
Calculated line absorption IR (FTIR) spectrum (A and C) and absorptive 2D IR spectrum (B and D) of amide I modes of R domain in unphos- and phos-CFTR in parallel polarization with population time t2=0. (E) Different spectrum generated by subtraction of unphos-CFTR from phos-CFTR 2D IR spectrum. In the 2D IR spectrum, dashed contour lines are negative (GB and SE signals), full contour lines are positive (EA absorption), the colored contour lines reflect equidistant intensity levels. In the FTIR spectrum, negative represent absorption.

Each absorptive 2D IR spectrum (Fig. 3B and D) is mainly composed of a doublet of peaks with opposite signs. The negative sign is an on-diagonal peak at ω_3_=ω_01_ due to the ground state to the first excited state (|0⟩−|1⟩) GB and SE signals of the sum of all the amide I modes in R domain, the EA absorption signals from the first excited state to the second excited state (|1⟩−|2⟩) appear at ω_3_=ω_12_ with the positive sign. Due to the anharmonic shift (∼16 cm^-1^) being smaller than the bandwidth of the transitions, the |0⟩−|1⟩ and |1⟩−|2⟩ transitions overlap and partially cancel. This results in negative peaks roughly distributed in the lower right corner of the diagonal, while positive peaks in the upper left corner. The overall shape of the 2D IR spectra of R domain in unphos- and phos-CFTR is almost identical, with the main difference being in the intensity, which is proportional to the square of the emitted electric field strength.

We made a difference 2D IR spectrum (Fig. 3E) generated by subtracting 2D IR spectrum of unphos-CFTR (Fig. 3B) from that of phos-CFTR (Fig. 3D) to clarify these intensity differences. The difference spectrum forms a pair of opposite peaks (− +) at the EA position (upper left corner of the diagonal) and a pair of opposite peaks (+ −) at the GB and SE position (lower right corner), where the two positive peaks form a single peak on the diagonal due to their close positions. This indicates that compared to unphos-CFTR, the absorption of phos-CFTR is weakened in the low-frequency band but enhanced in the high-frequency band. According to the emitted electric field *E*_*sig*_^*(3)*^ formulation in the method section, this means that phosphorylation may lead to an increase in the number of high-frequency oscillators or an enhancement of their transition dipole moments, while the opposite effect on the low-frequency oscillators.

### 4. Cross GB IR spectra of R domain in the unphos- and phos-CFTR

The 2D IR spectra with three types of Liouville space paths: GB, SE, and EA, although containing complete energy level transition information, peaks overlapping with multiple vibrational modes are especially problematic because it complicates the interpretation of spectra. This spectrum can be simplified by calculating the GB pathway on the oscillator transition from |0⟩−|1⟩. It contains all the information of site energies and coupling of oscillators that we care (19, 25). The main difference between the GB IR and 2D IR spectrum is that it does not calculate transition from |1⟩−|2⟩, which contains the information of anharmonic shift *Δ* _*ii*_≡ *ω*_*i*_^*12*^ *-ω*_*i*_^*01*^ (17) for the diagonal peaks of oscillator *i*. In fact, in the NISE calculating method, this anharmonic shift is assigned a constant value to follow the harmonic approximation. Therefore, from a computational perspective, ignoring this anharmonic shift will not lose the substantive information of oscillators.

In addition, the non-rephasing cross 2D IR spectra exhibit the highest sensitivity in detecting conformational variation (26, 27). Therefore, we calculated the non-rephasing GB IR spectrum in cross polarization for both unphos- and phos-CFTR in Fig. 4. It should be noted that in a non-rephasing GB(2D) IR cross spectrum, the cross peaks also lie on the diagonal (17). In Fig. 4A, we can see that in the R domain of unphos-CFTR, couplings mainly occur between oscillators at peak 1635 cm^-1^ and 1704 cm^-1^. Compared with unphos-CFTR, the coupling low-frequency peak of phos-CFTR (Fig. 4B) blue shift ∼4 cm^-1^, with an increase in intensity. The coupling high-frequency peak splits into two peaks of 1707 cm^-1^ and 1739 cm^-1^, and the intensity of the newly generated 1739 cm^-1^ peak is higher than that of the 1707 cm^-1^ peak. It is evident that the phosphorylation of R domain leads to a change in its internal coupling state. To clarify the amide I modes corresponding to the coupled oscillator pairs in this spectrum, we have to resolve the individual vibrational modes by calculating their eigenfrequencies, shown in the next section.

**Fig. 4.**
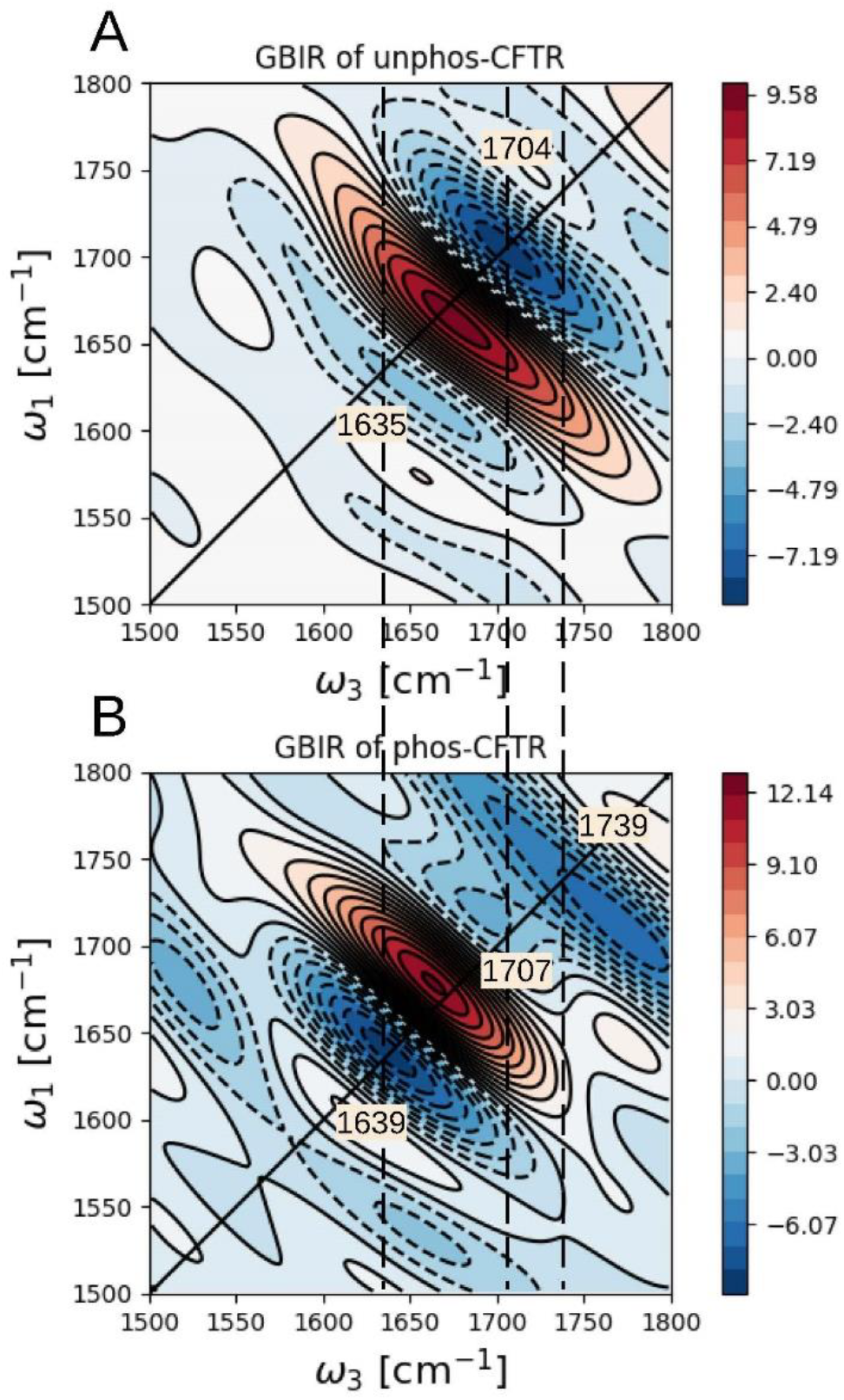
Calculated non-rephasing cross GB IR spectrum of amide I modes of R domain in unphos- and phos-CFTR with population time t2=0. The dashed contour lines are negative, full contour lines are positive. Unlike 2D IR spectrum, negative values represent absorptive contributions, while positive values represent the dispersive contributions from the R_GB_(ω_1_, ω_3_) response function. The colored contour lines reflect equidistant intensity levels.

### 5. Eigenstates of the oscillators

In a 3D structure of molecules, the local modes of oscillators will be coupled, so that they vibrate in unison. Regarding vibrational coupling, there are mainly two limiting cases to be discussed, namely strong coupling and weak coupling regime (17, 28). Taking coupled dimers as an example, when the coupling is small compared to the frequency splitting, |*β*_12_| ≪ |*ℏω*_2_ − *ℏω*_1_|, exciton states (vibrational excitons, the coupled local modes vibrate in unison) will be localized mainly on the individual sites. In the strong coupling regime, |*β*_12_| ≫ |*ℏω*_2_ − *ℏω*_1_|, exciton states will undergo delocalization.

We have used |∑_*j*_ *β*_*i*,*j*≠*i*_| and |*Δ*_*i*,*i±*1_| to replace the parameters on both sides of the relational inequality so as to extend it to the random coils in the R domain. Where |∑_*j*_ *β*_*i*,*j*≠*i*_| denote the coupling strength from all neighbors of oscillator *i*, |*Δ*_*i*,*i±*1_| is the frequency splitting of oscillator *i*. 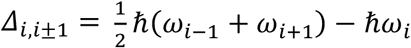 when *i* located in the middle of chain, *i* = 2,3, …, *N* − 1; *Δ*_*i*,*i±*1_ = *ℏω*_*i*_ − *ℏω*_*i±*1_ when *i* is at the end of the chain, *i=1 or N. ℏ* is the reduced Plank constant, *ω*_*i*_ denote the local vibrational frequency of oscillator *i, N* is the total number of oscillators.

We calculated the coupling strength |∑_*j*_ *β*_*i*,*j*≠*i*_| and frequency splitting |*Δ*_*i*,*i±*1_| of each oscillator in an average of 20 ps MD simulation (10000 frames) as shown in Fig. 5A. The results indicate that on the whole, the coupling strength is small than the frequency splitting, the coupling scheme of R domain belongs to weak coupling, especially in the random coils region (aa 671-804). In the weak coupling regime, oscillators of both the random coils and α-helices have the eigenstate energy 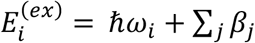 (17, 29). On this basis, we calculated the eigenfreqency of each exciton in an average of 20 ps trajectory and selected excitons located within ±5cm^-1^ of the cross peaks in Fig. 4 and recorded in Fig. 5B. We set the FWHM of the selected oscillator to be 10 cm^-1^ to match the natural linewidth (30). For unphos-CFTR, 5 oscillators located in the low-frequency domain, 24 oscillators located in the high-frequency domain. For phos-CFTR, 2 oscillators located in the low-frequency domain, 2 in the high-frequency domain.

**Fig. 5.**
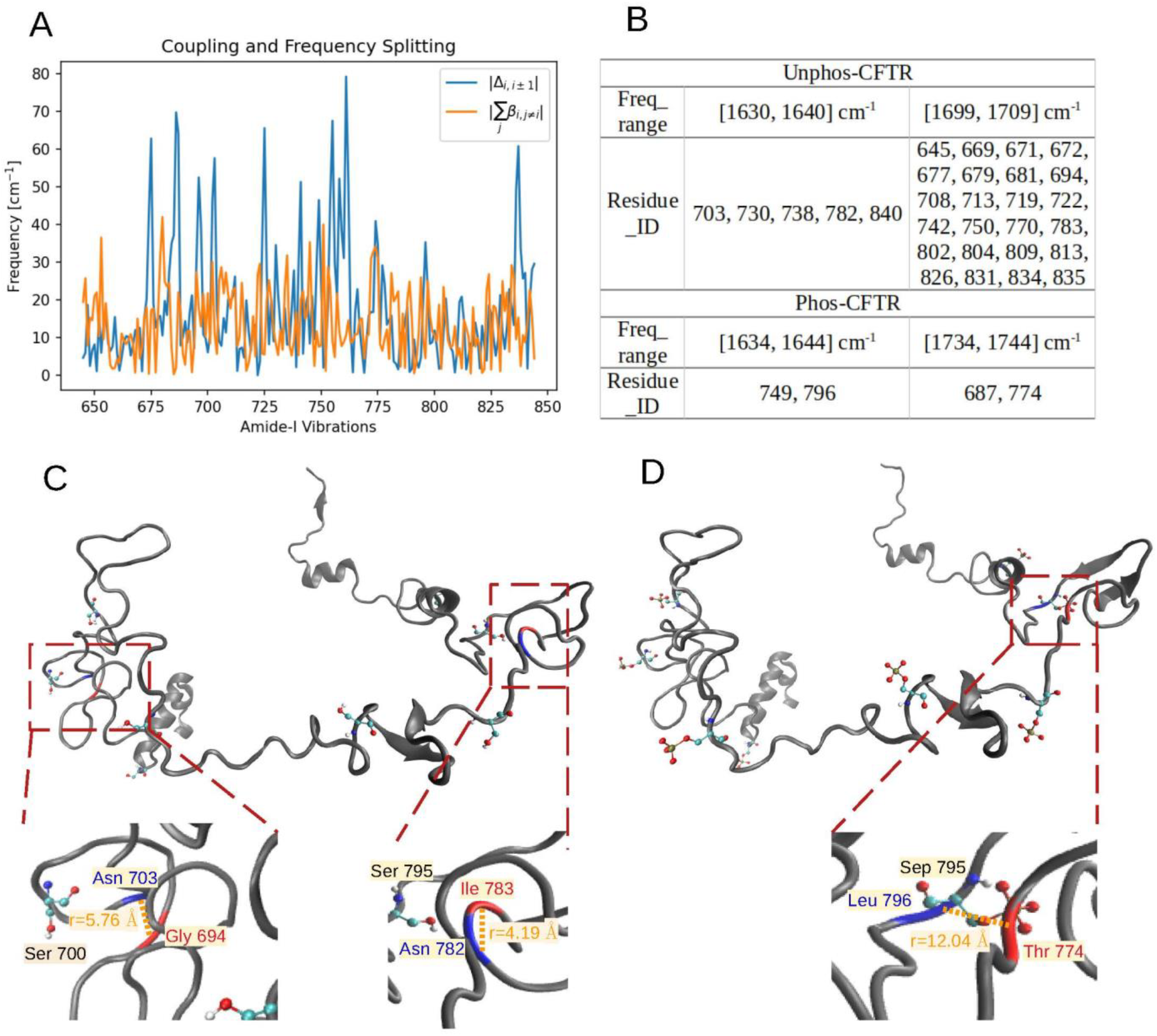
(A) The coupling and frequency splitting of each amide I mode in the R domain of phos-CFTR. (B) The amide groups of phos- and unphos-CFTR located within the FWHM of cross peaks in the non-rephasing cross GB IR spectrum. Here, “Residue ID” represent the amino acid residue number of R domain in CFTR. (C and D) The 3D structure of R domain in unphos- and phos-CFTR, respectively. The enlarged image of the red dashed boxes identify the coupled amino acid sites, where the second structures colored in blue represent low-frequency sites and red represent high-frequency sites, the corresponding amino acid residues are represented in situ by “amino acid name residue ID” and highlighted in relative colors. The distance between carbonyl bonds is represent by “r=xx Å” and highlighted with yellow backgrounds. The Ball-and-stick modes annotate the phosphorylation sites.

Then, we located these candidate oscillators from Fig. 5B in the 3D structure of unphos- and phos-CFTR, and screened those who may meet the basic coupling condition: distance ≤ 1.4nm, which is the cutoff distance of van der Waals and electrostatic force. We ultimately selected two amide I groups that meet the coupling condition in unphos-CFTR, as shown in Fig. 5C. Group 1 consists of Asn 703 and Gly 694, with a carbonyl bond distance of 5.76 Å, where we use the average distance between oxygen atom pairs and carbon atom pairs as the distance between the carbonyl bonds. Group 2 includes Asn 782 and Ile 783, with a distance of 4.19 Å. We selected one group in phos-CFTR: Leu 796 and Thr 774, with a distance of 12.04 Å (Fig. 5D).

Among all the phosphorylation sites, Ser 700 and Ser 795 are closest to the coupling oscillators mentioned above. To verify the effect of these couplings on molecular dynamics of CFTR, we only phosphorylated these two sites and conducted a 50 ns MD simulation in the same way as phos-CFTR. The corresponding kinetic parameters and GB IR spectra were calculated in Fig. 6. It is shown in Fig. 6A that in addition to the R domain, the fluctuations of C_α_ in the phos-700&795 CFTR are almost the same as those of phos-CFTR. Fig. 6C shows that within 50 ns MD simulation, the distance between the weighted COM of NBDs has the same trend as that of phos-CFTR. On the GB IR spectra (Fig. 6B and D), phos-700&795 CFTR and phos-CFTR have two similar cross peaks, with frequencies of 1645 cm^-1^ and 1745 cm^-1^, a blue shift ∼6 cm^-1^ compared to phos-CFTR. These results consistently confirm the corresponding relationship between phosphorylation, vibrational couplings, and the regulatory function of R domain.

**Fig. 6.**
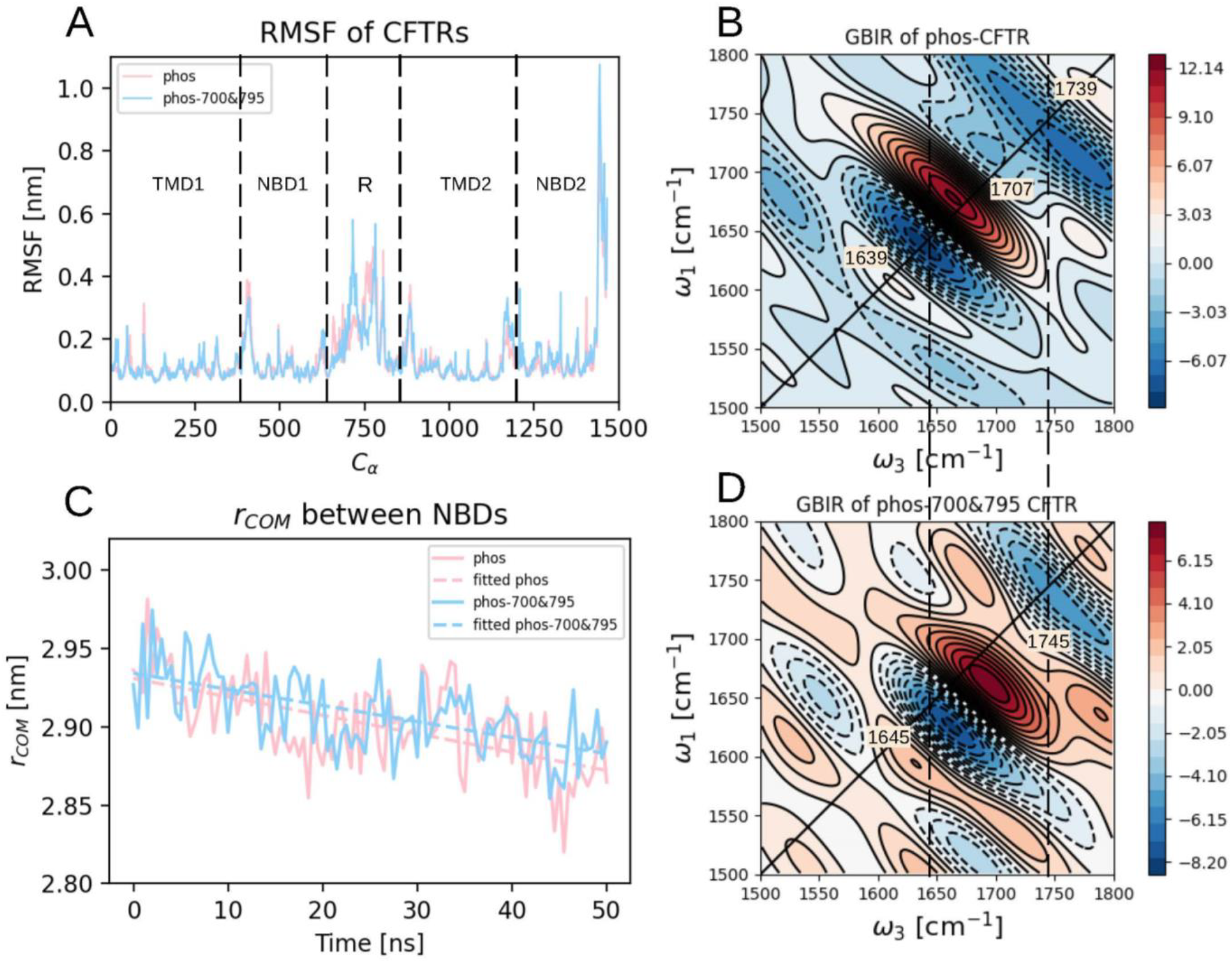
(A) The RMSF of Cα position in phos- and phos-700&795 CFTR in the trajectory of 50 ns after fitting to the initial frame. (C) The distance between the weighted COM of NBDs within 50 ns MD simulation. (B and D) Calculated non-rephasing cross GB IR spectrum of amide I modes of phos- and phos-700&795 CFTR with population time t2=0.

## Discussion

In this study, we utilized a combination of MD simulations and theoretically calculated non-rephasing cross GB IR spectra to characterize the vibrational couplings regulated by specific phosphorylation sites in the R domain of CFTR. Our MD simulations of unphos- and phos-CFTR indicate that phosphorylation of the R domain leads to the proximity of the two NBDs even without ATP binding. The non-rephasing cross GB IR spectrum of amide I modes in the R domain of unphos- and phos-CFTRs exhibit different cross peaks. By assigning these cross peaks to the eigenstates of specific excitons, we extracted two phosphorylation sites Ser 700 and Ser 795, which correspond to the kinetics of CFTR. Both the molecular dynamics characteristics of phos-700&795 CFTR and the vibrational coupling modes simulated by non-rephasing cross GB IR spectrum show consistent results with phos-CFTR, indicating that couplings between amide I modes regulated by phosphorylation of specific sites in R domain play a crucial role in regulating the kinetics of CFTR.

It has been established that opening of a CFTR channel is strictly coupled to the formation of a closed NBD dimer (31, 32). Other works (3, 10) have shown that the NBDs dimerized and TMD moved essentially as rigid bodies upon phosphorylation and ATP binding. Due to the easier quantification of the distance between NBDs compared to TMDs, we used the COM distance of NBDs as a quantitative criterion to evaluate the effect of phosphorylation on the conformational changes of CFTR, and used it in the following experiments to determine whether phosphorylation at specific sites can fully exert its function. We have confirmed through MD simulations that phosphorylation of R domain, which does not require ATP binding, can also lead to the proximity of the NBDs. Although the non-hydrolytic pathway of CFTR has been mentioned in other works (8, 33), referring to its ATP binding cassette (ABC) (34) structural characteristics, we tend to believe that phosphorylation is a initiating factor for NBDs dimerization. In addition, the works that claim basal phosphorylation of some CFTR residues are essential for subsequent full channel activation by PKA (35, 36) support our speculation as well.

The conventional site-directed mutagenesis (37) and the caged-serine method (38) have suggested a degree of non-equivalency among sites, with Ser 813, 795, 660, 670, 700 inferred to be stimulatory, but Ser 768 to be inhibitory, and Ser 737 to be debated. Consistent with the above results, our study confirms the stimulatory effect of Ser 700 and 795, and confirms that this effect originates from the changes in the vibrational coupling state of neighboring amide I modes after phosphorylation of these sites.

The MD equilibrium random coils of R domain is located on the surface of CFTR, with N-terminal α-helices embedded in the TMD-NBD halves. This semi-embedded structure can simultaneously explain the electron density of R domain docked in the intracellular vestibule of the cryo-EM structure (10), as well as how PKA efficiently contacts its interaction sites. In addition, we have also found that the conformation of the R domain did not undergo significant changes before and after phosphorylation, nor was it released from the TMD-NBD halves as predicted (3). This may be influenced by our experimental setup, which does not include the role of ATP. Therefore, there is not enough energy to promote the aggregation of NBDs and the release of R domain.

The time dependence of the Hamiltonian resulted from the fluctuations of solvent and molecular structure, is the source of infrared lineshape, and can also cause energy transfer between eigenstates that will alter the intensity of cross and diagonal peaks or create completely new peaks (39). From Fig. S3, it can be seen that with the delay of population time t_2_, phos-CFTR exhibits the fastest population relaxation, which is suggested to be via the population transfer from amide I mode to amide II mode (CN stretch and NH bend) and hydrogen-bond vibrations (40). Therefore, phosphorylation of the R domain may lead to faster population transfer from amide I modes, further accelerating the release of existing coupling modes in R domain.

By calculating the eigenfrequency of excitons and matching them with the cross GB IR spectrum, we can locate the intermodal couplings with the individual vibrational modes. Although in this paper we used the non-rephasing cross GB IR spectrum, which has higher resolution than traditional 2D IR spectrum, theoretically, all IR spectra (including FTIR and 2D IR spectra) are applicable to the above method. This provides an implementation approach for achieving bond-specific structural resolution to resolve the dynamics of protein skeletons, which can be used to study the structural and functional characteristics of proteins at the biophysical level.

## Methods

### 1. MD Simulations

MD simulations were performed with GROMACS (41) packages using the gromos43a1p-4.5.1.ff force field translated by Justin Lemkul and Graham Smith, which was extended to include phosphorylated residue with the parameter set by Tomas Hansson (42, 43). The primary CFTR structure (PDB ID: AF_AFP13569F1) has a full sequence length of 1480, it is a predicted AlphaFold model. MD simulations have been performed on this model in a solvated lipid membrane environment to obtain an equilibrium subprime structure with a folded R domain. This subprime structure will be used as the initial structure for all the remaining simulations.

For all MD simulations, proteins were terminated with NH_3_^+^ and COO^-^ on the N- and C-terminus. All bonds (even heavy atom – H bonds) are constrained with LINCS algorithm, allowing an integration time step of 2 fs. The long-range electrostatic interactions were calculated by the particle mesh Ewald (PME) method with a grid spacing of 0.16 nm. The short range repulsive and attractive dispersion interactions were described using the Lennard-Jones potentials, with a cutoff length of 1.4 nm. The temperature of the system was kept constant by the Nosé-Hoover coupling method on Protein_DOPC and Water_and_ions separately with external heat bath at 310 K and a coupling time of 0.5 ps. The pressure was maintained constant through the Parrinello-Rahman coupling method, with a pressure bath of 1.01325 bar and a coupling time of 2.0 ps. The Convergence standards and errors were set by software default.

Phosphorylation was achieved by replacing the hydroxyl hydrogen on the serine side chain with phosphite (-PO_3_^2-^). The DOPC lipid bilayer (44) was used to simulate the cytoplasmic membrane environment, and in our work, it has been expanded to a system with 1152 lipids and 43425 water molecules. A 20 ns simulation at 310 K has been used to stabilize this expanded system. 43A1-S3 force field (45) was used for lipids, and the SPC model is used for water molecules.

The Lambada and Inflategro2 (46) method were used to embed the protein into the DOPC lipid bilayers, and solvate with water. Chlorine ions have been added to maintain the electrical neutrality for this system, the solvated system is shown in Fig. S1.

### 2. 2D IR Spectra Simulations

For each configuration, a 20 ps MD simulation was performed to store the structure every 2 fs, giving a trajectory with a total of 10000 snapshots. For each trajectory, the AIM program (47) was used to generate a time-dependent Hamiltonian and transition dipole moments of each amide I mode in the R domain. The site frequencies were generated using a Jansen map approach, which utilizes the electrical field and its gradient on the C, N, O, H atoms of amide bonds. Frequency correction is achieved by using a residue-type-depended mapping based on the Ramachandran angles (48) of the bond. The anharmonicity needed for describing doubly excited states was set to 16 cm^-1^ for all units. The couplings between both the nearest and not nearest neighbor amide groups were calculated using the transition charge coupling (TCC) scheme (49), which determines the interactions between any pair of atoms inside the amide groups. In the spectral simulation, the side-chain carbonyl vibrations were neglected.

The numerical integration of the Schrödinger equation (NISE) method (19) was used for calculating both the linear absorption spectra and the 2D IR spectra. The linear absorption spectrum is obtained by Fourier transform of first-order response function. 2D IR spectrum in frequency domain is obtained by a two-dimensional Fourier transform of third-order response function, where the first time delay *t*_*1*_ gives the *ω*_*1*_ frequency axis, the last time delay *t*_*3*_ gives the *ω*_*3*_ frequency axis of spectrum. 2D IR spectroscopy is a four wave mixing technique (17, 50), the spectrum is given as the sum of the signal emitted in the directions with wave vectors 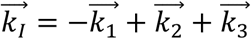 and 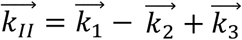, where 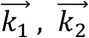 and 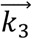 are the wave vectors of the incoming infrared fields. 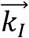 is the rephasing signal and 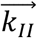 is the non-rephasing signal. Each of these signals contains three contributions from distinct Liouville space pathway: ground-state bleach (GB), stimulated emission (SE), and excited-state absorption (EA) (25). The emitted electric field *E*_*sig*_^*(3)*^ of 2D IR spectroscopy is proportional to the third-order response function *R*^*(3)*^, oscillator density *N*, vibrational frequency *ω*, and the electric field strength *E* of each of the three laser pulses (17), as 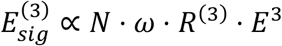 . As the third-order response function *R*^*(3)*^ is proportion to the 4th power of the transition dipole moment *μ* of oscillators, for the systems we constructed with the same incoming electric field strength, the emitted field can be simplified as 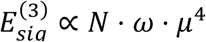, which fluctuates with the transition dipole, vibrational frequency of the oscillators and their density.

## Acknowledgments

This work is supported by grants from the National Nature Science Foundation of China (Grant No. 82173388 and No. 22102163). We also appreciate the software support provided by Thomas la Cour Jansen.

