## Supplementary material for "Probing Phosphorylation-Induced Vibrational Couplings in CFTR by 2D IR Spectra Simulations": Fig. S1

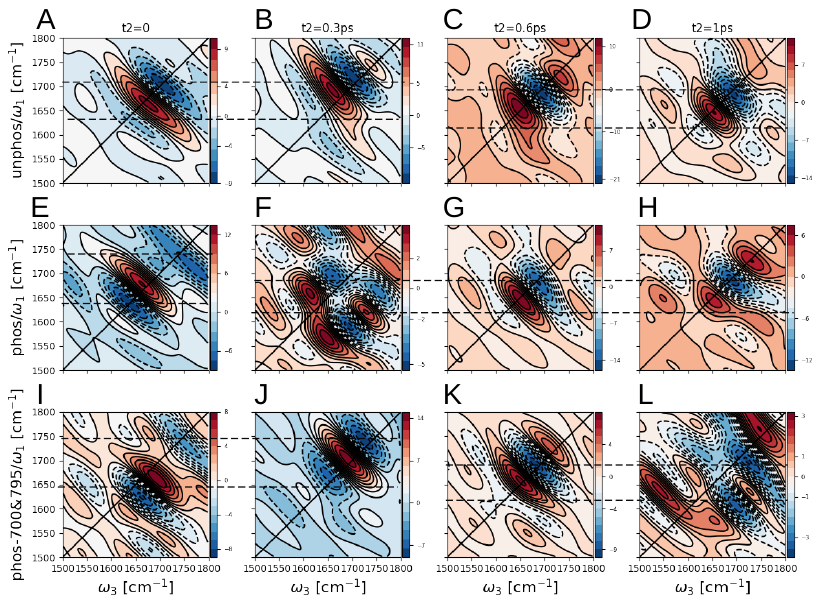

**Fig. S3.** Calculated non-rephasing cross GB IR spectrum of amide I oscillators of unphos-, phos-, and phos-700&795 CFTR in cross polarization with population time t2=0, t2=0.3 ps, t2=0.6 ps, t2=1 ps. In the GB IR spectrum, dashed contour lines are negative, full contour lines are positive as well. However, in different from the 2D IR spectrum, negative represent the absorptive contributions, positive represent the dispersive contributions from the R_GB_(ω_1_, ω_3_) response function. The colored contour lines reflect equidistant intensity levels.
