## Supplementary material for "Probing Phosphorylation-Induced Vibrational Couplings in CFTR by 2D IR Spectra Simulations": Fig. S2

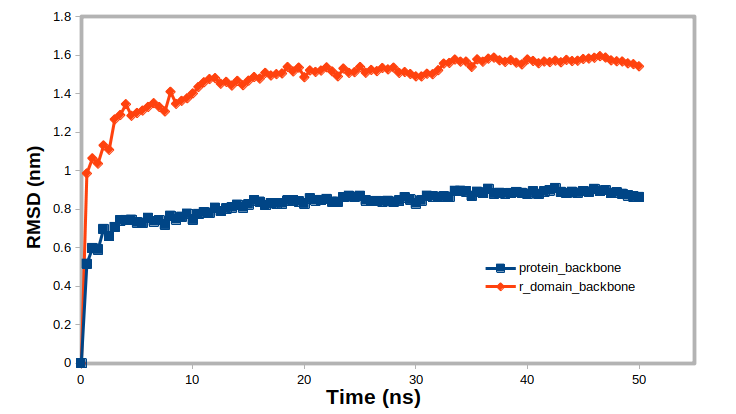

Fig. S2. The root mean square deviation (RMSD) of the backbones of whole protein and the backbones of the R domain from the 50ns MD trajectory to the first frame.
