## Supplementary material for "Probing Phosphorylation-Induced Vibrational Couplings in CFTR by 2D IR Spectra Simulations": Fig. S3

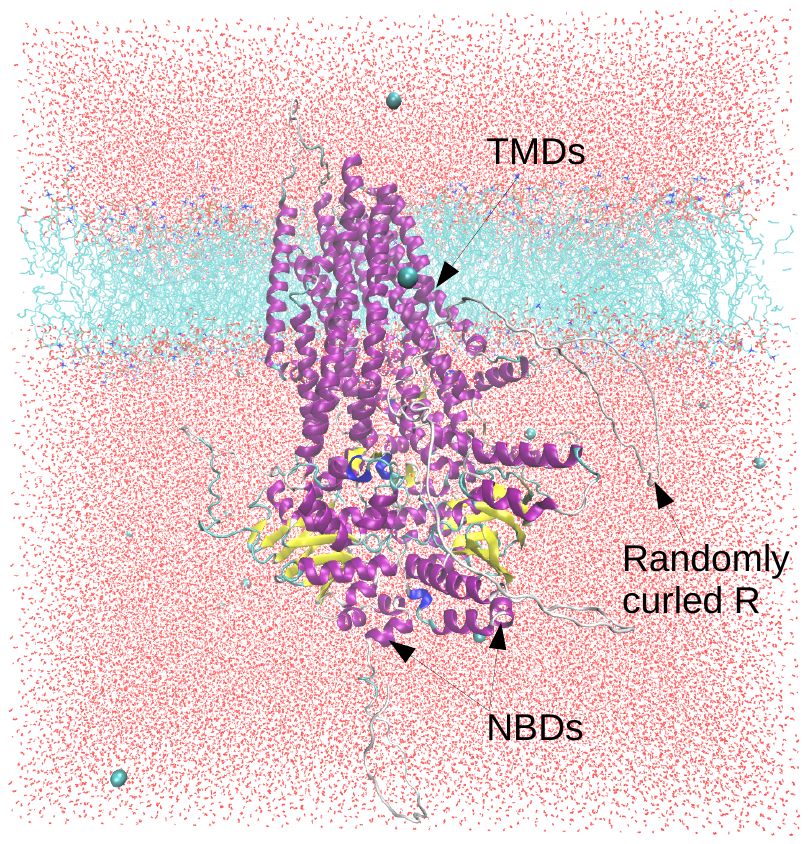

Fig. S1. The solvated box. The CFTR model predicted by AlphaFold is embedded in the DOPC lipid bilayers, and surrounded by SPC water and chloride ions. Box size: 20nm × 20nm × 20nm.
